# Effect of mechanical stretching stimulation on maturation of human iPS cell-derived cardiomyocytes co-cultured with human gingival fibroblasts

**DOI:** 10.1101/2023.12.15.567696

**Authors:** Mengxue Wang, Harumi Idei, Chen Wang, Yin Liang, Yun Liu, Yusuke Matsuda, Ken Takahashi, Hiroshi Kamioka, Keiji Naruse

## Abstract

In the realm of regenerative medicine, despite the various techniques available for inducing the differentiation of induced pluripotent stem (iPS) cells into cardiomyocytes, there remains a need to enhance the efficiency of this induction process. This study aimed to improve the differentiation efficiency of iPS-derived cardiomyocytes (iPS-CMs) by incorporating mechanical stretching. Human iPS cells were co-cultured with human gingival fibroblasts (HGF) on a polydimethylsiloxane (PDMS) stretch chamber, where mechanical stretching stimulation was applied during the induction of cardiomyocyte differentiation. The maturation of iPS-CMs was assessed using qRT-PCR, immunofluorescence staining, transmission electron microscopy, and contractility comparisons. Results indicated significantly elevated gene expression levels of cardiomyocyte markers (cTnT) and the mesodermal marker (Nkx2.5) in the stretch group compared to the control group. Fluorescent immunocytochemical staining revealed the presence of cardiac marker proteins (cTnT and HCN4) in both groups, with higher protein expression in the stretch group. Additionally, sarcomere length in the stretch group was notably larger than in the control group. A significant increase in the contractility of iPS-CMs was observed in the stretch group. These findings demonstrate that mechanical stretching stimulation enhances the maturity and differentiation efficiency of iPS-CMs co-cultured with fibroblasts.

## 1. Introduction

Regenerative medicine using induced pluripotent stem (iPS) cells has been studied in various fields in recent years, and has already been clinically applied in some areas such as the retina ^1^. In the field of heart, research has been developed e.g., treatment with iPS cell-derived myocardial sheets ^2^, and autologous heart transplantation by creating a heart from iPS cells is also expected to have great clinical application in the future field.

Heart transplantation is the only treatment for end-stage heart failure that has been difficult to treat with pharmacotherapy and auxiliary circulation alone. At the same time, resource of heart transplantation is limited and are available for only a limited number of patients. If it becomes possible to create a heart with iPS cells, this problem of waiting for transplantation can be solved at once. However, although various methods have been devised to induce differentiation of iPS cells into myocardial tissue, and a large amount of myocardial tissue has been obtained ^3^. In terms of clinical application, not only should differentiation induction efficiency be maintained, but also the myocardial tissue maturity should be retained ^4^.

Although cardiomyocytes compose 75% of the volume of normal myocardial tissue, they account for only 30%-40% of the total number of cardiac cells. Most of the remaining cells are non-myocytes, mainly fibroblasts ^5^. Many researches have been conducted to promote myocardial maturation by simulating the cardiac environment. Ieda et al. found that mice cardiomyocytes (CMs) co-culture with cardiac fibroblasts (CFs) can induce maturation through cell-to-cell interactions and paracrine signaling ^6^.Co-culture with mouse embryonic fibroblasts improves the differentiated phenotype of mouse embryonic stem cell-derived CMs ^7^. Also, human pluripotent stem cell-derived CMs co-cultured with human foreskin fibroblasts promote maturation of cardiac tissue by supplementation with ascorbic acid and increasing static stretching ^8^. Recently, Giacomelli et al. showed the combination of human iPS cells-derived CMs, CFs and cardiac endothelial cells (ECs) promote CMs maturation, and inclusion of iPS-CFs further enhanced structural, electrical, mechanical and metabolic maturation of CMs ^9^.

The previous studies have suggested that fibroblasts are factors that promote iPS cell differentiation and influence myocardial tissue maturation. Matsuda et al. demonstrated that co-culture with human gingival fibroblasts (HGF) facilitates differentiation of iPS cells into cardiomyocytes (iPS-CMs) with higher contractility ^10^. It has also been reported that mechanical stretching stimulation promotes the maturation of iPS cell-derived cardiac tissue ^11^. HGF are easy to obtain during routine dental procedures and the wound heals quickly with very little scarring. As an easily accessible cell source, HGF has progenitor/stem cell-like differentiation capabilities for tissue engineering and regenerative applications ^12^.

Therefore, in this study we aimed at improving the differentiation induction efficiency of iPS-HGF co-culture into myocardial tissue by applying mechanical stretching. The maturity of the iPS-CMs was evaluated by the level of gene and protein expression, sarcomere length and contractility. These results could provide potential cell model in the progression for cardiac maturation study.

## 2. Materials and methods

### 2.1. Isolation of HGF

When extracting teeth from a patient undergoing orthodontic treatment at Okayama University Hospital, about 2 mm^3^ of gingival tissue was collected (Figure 1B). Patients were at least 16 years of age. Patients with periodontitis were excluded for tissue collection. The tissue pieces were then minced in a tissue culture dish with the following solution: 10% fetal bovine serum (FBS: Sigma Aldrich, MO, USA), 1 mol / l-HEPES buffer solution (Nacalai Tesque, Kyoto, Japan), penicillin-streptomycin, and Dulbecco’s Modified Eagle Low Glucose Medium (Thermo Fisher Scientific, MA, USA). Finally, the cells were incubated at 37°C in a 5% CO_2_ humidified incubator. HGF that became confluent in the tissue culture dish was periodically subjected to trypsin treatment and subcultured. All experiments were performed with HGF between passages 3 and 10. All HGF handling and experimental procedures in this study were approved by the Ethics Committee of the Graduate School of Biomedical Sciences, Okayama University (approval number 1612-007-002). Informed consent was obtained from the subject.

**Figure 1.**
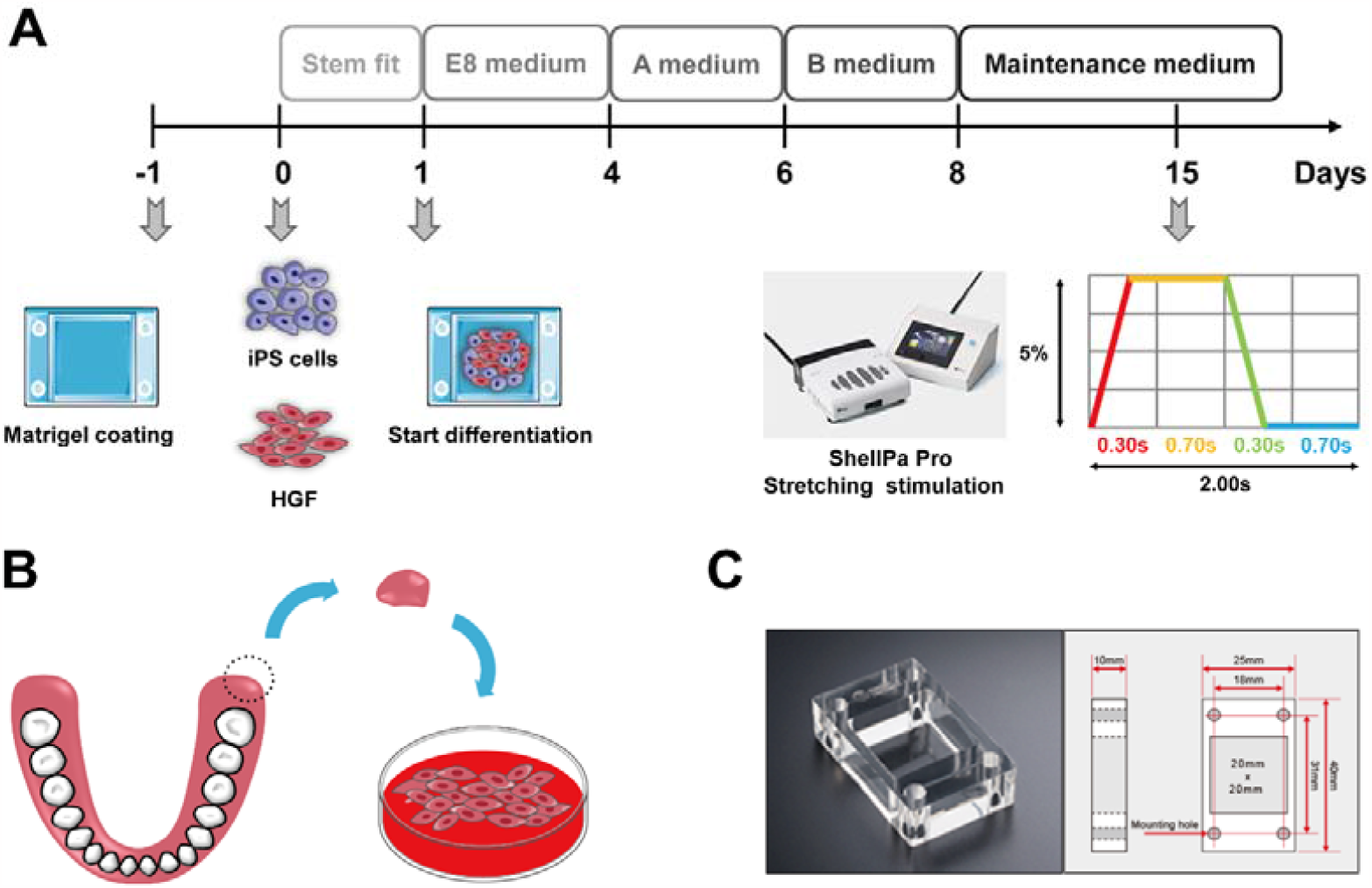
Protocols for myocardial differentiation of iPS cells co-cultured with HGF and mechanical stretching stimulation system. **A** Myocardial differentiation protocol using iPS cells co-cultured with HGF. **B** Protocol for the collection of human gingival tissue and isolation of HGF. **C** Polydimethylsiloxane (PDMS) stretch chamber.

### 2.2. Culture of iPS cells

Human iPS cells (type 201B7) were purchased from RIKEN (Kyoto, Japan). StemFit AK02N medium (Ajinomoto, Tokyo, Japan) was used for maintenance culture of iPS cells. A 6-well plate was coated with laminin 511-E8 fragment diluted to 0.5 g / ml in PBS (2 ml per well), and the plate was allowed to stand overnight at 4°C. Then the laminin 511-E8 fragment solution was removed from the culture dish of iPS cells, and the iPS cells were washed with PBS. Subsequently, 800 µl of TrypLE Select (Life Technologies, Carlsbad, CA, USA) was added to the wells and incubated for 7 minutes in a humidified incubator at 37°C, 5% CO_2_. Next, TrypLE Select was removed, the cells were washed again with PBS, and 1 ml of StemFit AK02N medium containing inhibitor of Rho-associated, coiled-coil containing protein kinase, Y-27632 (10µM) was added. Next, the iPS cells adhered to the bottom of the culture dish were peeled off using a scraper, and seeded at a density of 3.0×10^4^ cells/well on a 6-well culture plate coated with laminin 511-E8 fragment. The next day, the medium was removed and replaced with StemFit AK02N medium without Y-27632. The medium was changed on days 1, 4, 5, and 6. All experiments were performed with iPS cells between passages 7 and 20.

### 2.3. Induction of differentiation of iPS cells into myocardial tissue

All samples were cultured using a PDMS (polydimethylsiloxane) stretch chamber (size: 20×20 mm, Menicon, Aichi, Japan) to apply mechanical stretching stimulation during the induction of differentiation into cardiomyocytes (Figure 1A and C). First, Matrigel (Corning, NY, USA) was diluted to 35.5µl/ml in Dulbecco’s Modified Eagle Low Glucose Medium (Thermo Fisher Scientific, MA, USA). 1.4 ml/well Matrigel solution was added to wells, and the plate was kept overnight at 4°C for coating. The next day, 4.9×10^5^ cells of iPS cells and 2.1×10^5^ cells of HGF were mixed and seeded on the wells. From the day after the seeding, the cells were cultured in Essential 8 medium (Thermo Fisher Scientific, MA, USA) for 3 days, and the medium was changed every day. Subsequently, differentiation induction of iPS cells was started using PSC cardiomyocytes Differentiation Kits (Thermo Fisher Scientific, MA, USA) according to the manufacturer’s protocol. Briefly, the medium was removed and cardiomyocytes differentiation medium A was added. Two days later, the medium was removed, and cardiomyocytes differentiation medium B was added. After incubation for 2 days, the medium was removed and cardiomyocytes maintenance medium was added. The cardiomyocytes maintenance medium was changed every other day thereafter. On the 15th day from the start of the culture, a mechanical stretching stimulus (stretching rate: 5%, frequency: 0.5Hz) was applied for 72 hours using ShellPa Pro stretching equipment (Menicon, Aichi, Japan) (Figure 1A). A sample cultured in a stationary state without applying a stretching stimulus was used as a control group.

### 2.4. Quantitative RT-PCR

RNA was extracted using the High Pure RNA Isolation Kit (Roche, IN, USA) and reverse transcribed into cDNA using the Verso cDNA synthesis kit (Thermo Fisher Scientific, MA, USA). The target genes were cardiac marker cTnT, mesodermal marker Nkx2.5, pluripotent marker Sox2 and Oct4, and the endogenous control was 18S rRNA. The primers of these genes were mixed with SYBR green reagent (Life Technologies, Warrington, UK) and quantitative RT-PCR was performed to quantify the expression level of the target genes. Table 1 shows the nucleotide sequences of the primers. The expression level of the target genes was normalized by the expression level of 18S rRNA, and calculated by the ΔΔCt method. All measurements were performed in triplicate for each target gene in three independent samples.

**Table 1.**
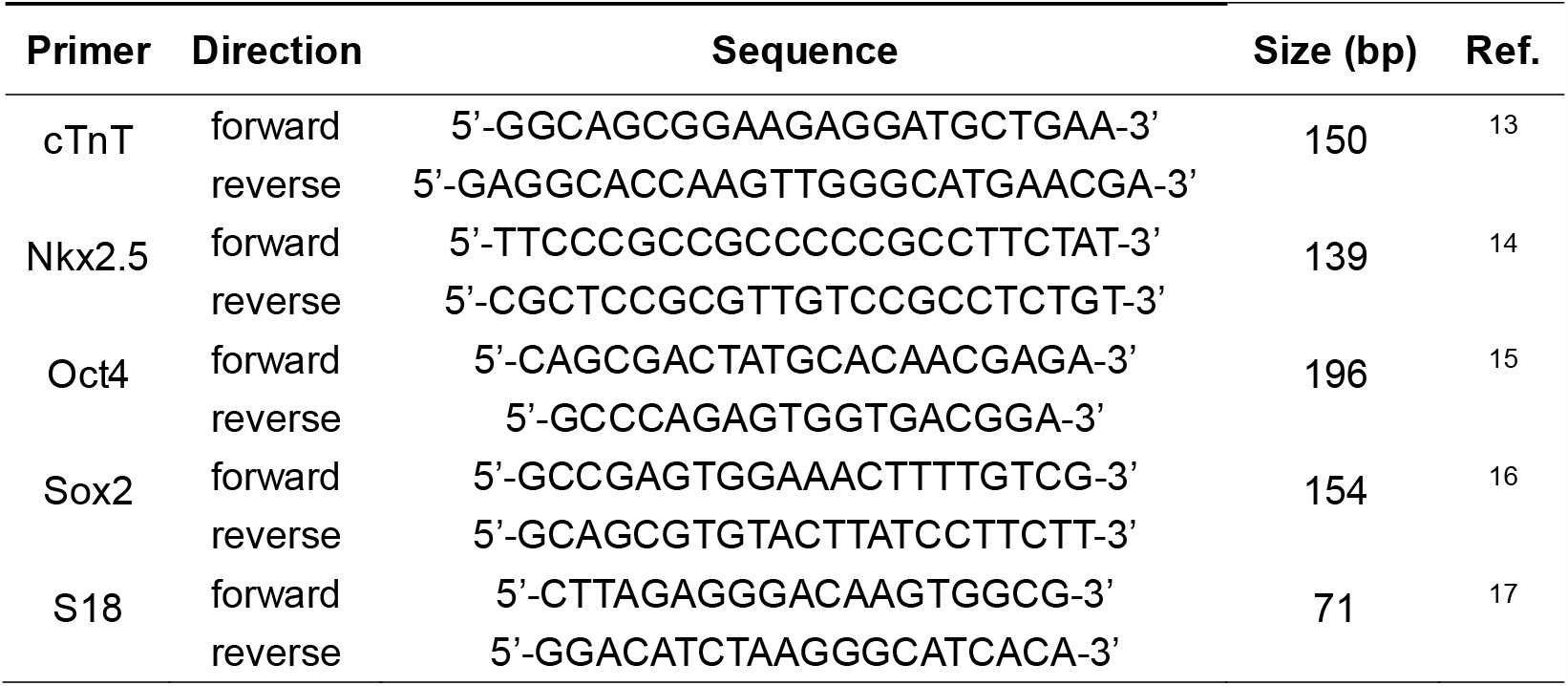
Primers used for quantitative RT-PCR.

### 2.5. Fluorescent immunocytochemistry

The expression of the cardiac muscle marker was evaluated by fluorescent immunocytochemical staining. Prior to staining, cells were fixed with 4% paraformaldehyde in PBS for 15 minutes at room temperature, followed by three times of PBS washing. Next, the cells were permeabilized with 0.2% Triton X-100 (Nacalai Tesque, Kyoto, Japan) for 15 minutes and blocked with 3% bovine serum albumin (BSA: Sigma Aldrich, MO, USA) for 30 minutes. Subsequently, the primary antibodies were added and cells were incubated at 4°C overnight. Next, after washing three times with PBS, the secondary antibodies were added and incubated at room temperature for 1 hour. Nuclei were stained by incubating with PBS supplemented with NucBlue reagent (Thermo Fisher Scientific, MA, USA) for 5 minutes. The following antibodies were used. Primary antibodies: cardiac troponin T mouse monoclonal antibodies (cTnT: Thermo Scientific, Rockford, IL, USA), diluted to 1/200 with 3% BSA, and hyperpolarization-activated cyclic nucleotide-gated channel 4 monoclonal antibody (SHG 1E5) (HCN4: Thermo Scientific, Rockford, IL, USA), diluted to 1/100 with 3% BSA. Secondary antibodies: goat anti-mouse antibodies Alexa Fluor 488 (Life Technologies, UK) and goat anti-rat antibody Alexa Fluor 568 (Life Technologies, UK), both were diluted 1/1000 with 3% BSA. Fluorescence images were acquired using a confocal laser microscope (objective lens: 10×, 40×, LSM780, Carl Zeiss, Oberkochen, Germany).

### 2.6. Transmission electron microscope observation

The cells were pre-fixed overnight at 4°C in a 0.1 M cacodylate buffer containing 2% glutaraldehyde and 2% paraformaldehyde. After washing with a 0.1 M cacodylate buffer, the cells were post-fixed with 2% OsO_4_ for 1.5 hours at 4°C, then washed with 0.1 M cacodylate buffer and dehydrated with ethanol. Then, they were embedded in Spurr resin (Polyscience) and thermally polymerized, and an ultra-thin section of 80 nm was prepared using ultramicrosome (LEICA EM UC7) (Leica Microsystems, Wetzlar, Germany). They were double stained with uranium and lead, and observed using a transmission electron microscope (H-7650, Hitachi High-Technologies, Tokyo, Japan). Image J software (US National Institute of Health, Bethesda, Maryland, USA) was used for sarcomere length measurement.

### 2.7. Video analysis of iPS-CMs

Videos of the contraction of the iPS-CMs were recorded for 10 seconds before and after mechanical stretching stimulation using a phase contrast microscope (BZ-X710, KEYENCE, Osaka, Japan). All videos were captured at a fixed position using a 4×objective lens at 20 frames per second. The displacement vector fields were calculated between frame 1 and all subsequent frames (frame 1 vs 2, frame 1 vs 3, frame 1 vs 4, etc.). The displacement vector fields were calculated by using the particle image measurement plug-in ^18^ of image J (National Institutes of Health ^19^), and the displacement vector D (x, y) was obtained for each 16×16 pixel. The maximum displacement vector M (x, y) was defined as follows for each (x, y) pair:

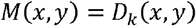

Here, k represents the frame number at which | D_k_ (x, y) | = max [| D_2_ (x, y) |, | D_3_ (x, y) |, …, | D_n_ (x, y) |]. The number n indicates the last frame number (n = 200 in this study). The contractility C was calculated in arbitrary units as follows:

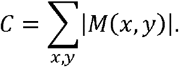

The rates of change in contractility before and after mechanical stretching stimulation were determined as follows:

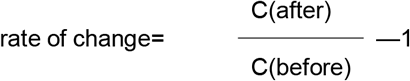

### 2.8. Statistical analysis

Data obtained from experiments were expressed as mean ± standard error of the mean (SEM). Unpaired t-test was used for comparison between the two groups. Differences were considered significant when p <0.05.

## 3. Results

### 3.1. Expression of myocardial markers and pluripotent markers iPS-CMs

To compare expression levels of differentiation marker genes in response to mechanical stimulation, we performed quantitative reverse transcription polymerase chain reaction (qRT-PCR). Cardiac troponin T (cTnT) was used as cardiomyocytes marker and Nkx 2.5 was used as a mesodermal marker. Expression level of cTnT was larger in the stretch group compared to control group (1.939 ± 0.350 and 4.818 ± 0.737, respectively). Similarly, mRNA expression of Nkx2.5 was significantly increased in the stretch group compared to control group (1.232 ± 0.147 and 2.601 ± 0.341, respectively) (Figure 2A). Therefore, these results suggested that mechanical stretching promotes the differentiation of iPS cells into cardiomyocytes. On the other hand, although expression level of pluripotent marker Oct4 tended to decrease in response to stretch compared to control group (0.340 ± 0.165 and 0.246 ± 0.105, respectively), but the difference was not statistically significant. Similarly, expression level of pluripotent marker Sox2 tended to decrease in response to stretch compared to control group (0.556 ± 0.186 and 0.177 ± 0.048, respectively), but the difference was not statistically significant (Figure 2B).

**Figure 2.**
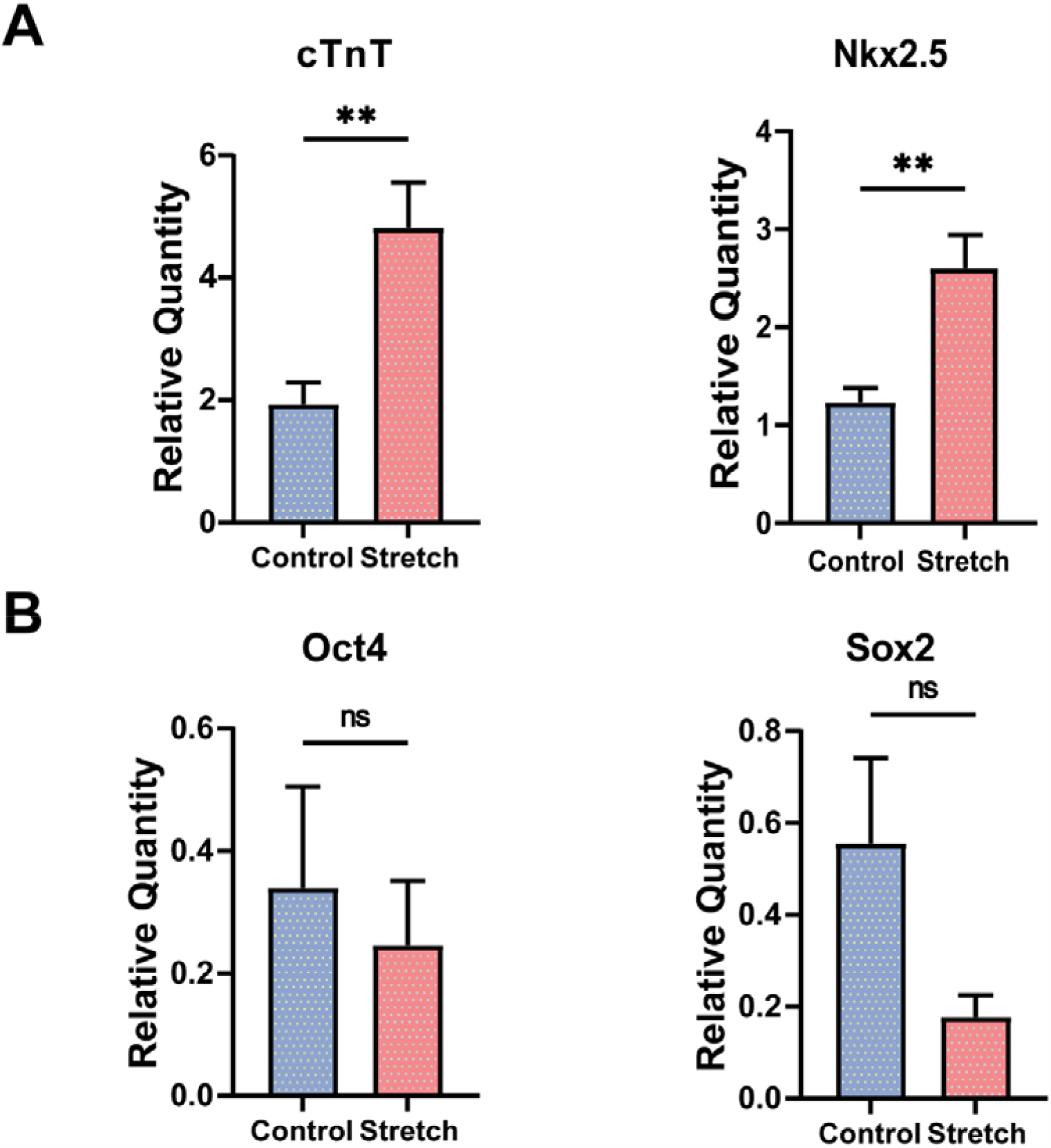
mRNA expression of cardiac and pluripotent markers. qRT-PCR was employed to analyze the mRNA expression levels of **A** cardiac markers (cTnT, Nkx2.5) and **B** pluripotent markers (Oct4, Sox2) in iPS-CMs between the control and stretch groups. The data are presented as means ± S.E.M. n = 5 for each condition. Statistical analysis was conducted using an unpaired t-test. ns: P > 0.05, **: P < 0.01.

### 3.2. Immunofluorescence staining of iPS-CMs

Next, we observed expression of cardiac marker proteins, cTnT and HCN4 by immunofluorescence. Expression of these cardiac marker proteins were evident in both stretch and control groups (Figure 3A). Striated pattern of cTnT was observed in the stretch group, implying developed sarcomere structure. The expression of cTnT and HCN4 was higher in the stretch group compared to control group (Figure 3B). Thus, these results indicated mechanical stretching promotes the maturation of cardiac tissue.

**Figure 3.**
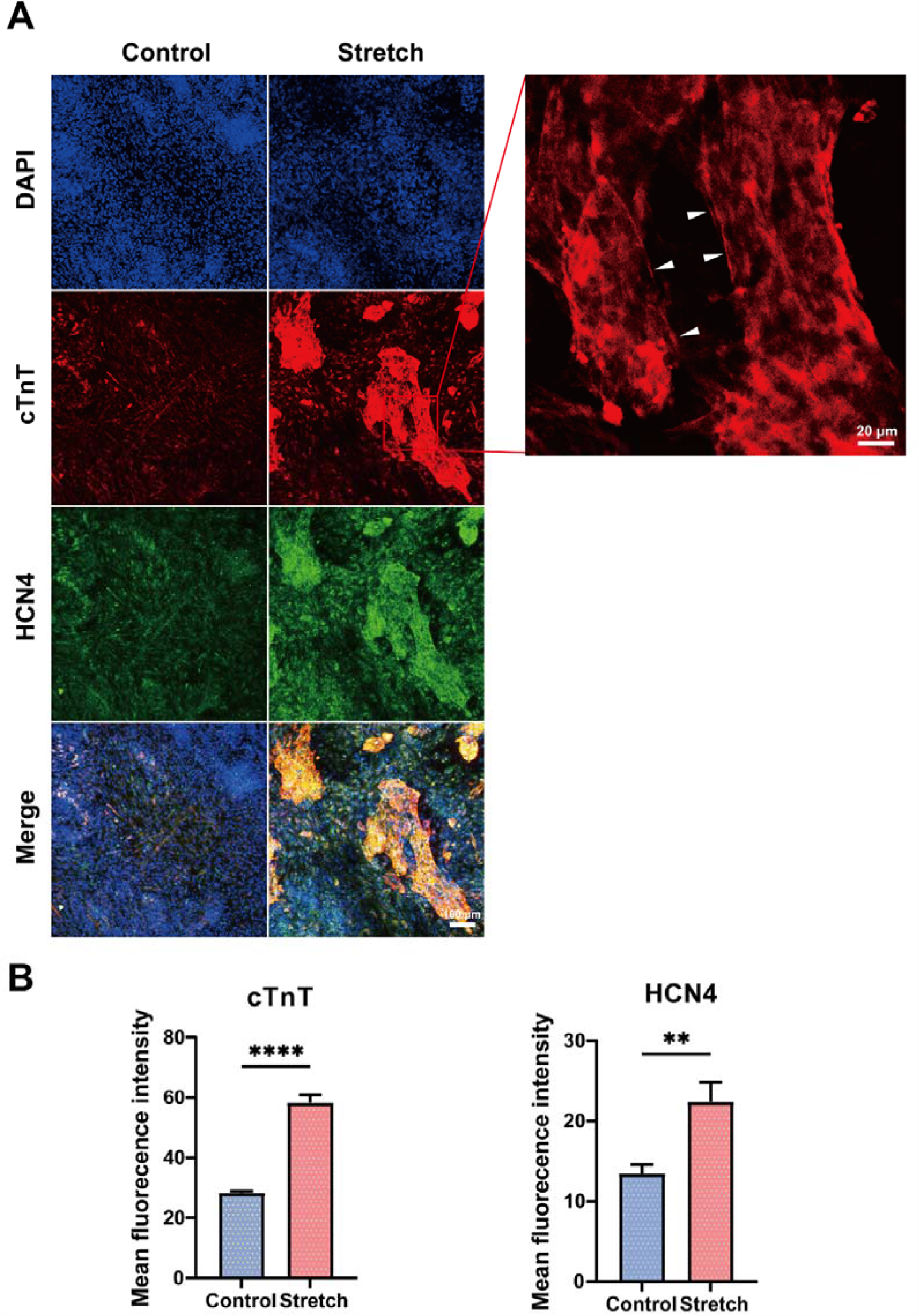
Protein expression of cardiac markers. **A** Immunofluorescence imaging of iPS-CMs in the control and stretch groups. Blue: DAPI, red: cTnT, green: HCN4. In the magnified image (right), arrowheads indicate sarcomere structure. **B** Mean fluorescence intensity of cTnT and HCN4 was compared between the control and stretch groups. The data was presented as means ± S.E.M. For the control group, n = 13; for the stretch group, n = 9. Statistical analysis was performed using an unpaired t-test. **: P < 0.01, ****: P < 0.0001.

### 3.3. Morphological comparison of microstructure of iPS-CMs by transmission electron microscope

In order to compare the structures of the iPS-CMs in the control group and stretch group in more details, observations were made with a transmission electron microscope (Figure 4A). A series of sarcomere structures were observed in both groups. While the sarcomere length in the control group was 1.232 ± 0.024 μm, that in the stretch group showed a significantly larger value of 1.537 ± 0.020 μm (Figure 4B). Besides, iPS-CMs in the stretch group showed a higher degree of alignment than in the control group. Therefore, the sarcomere structure can become more mature through mechanical stretching stimulation.

**Figure 4.**
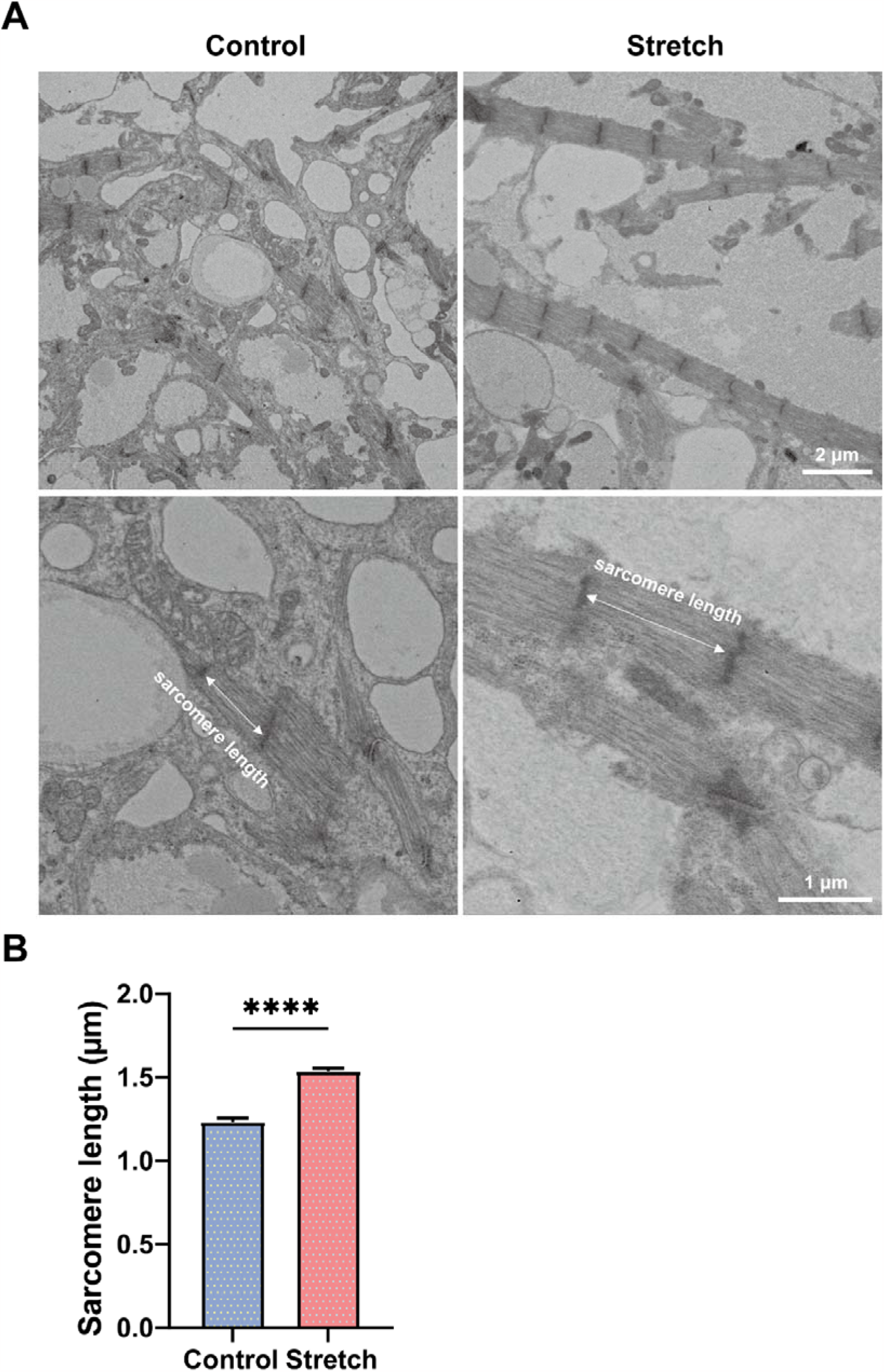
Transmission electron microscopy images of iPS-CMs. **A** Transmission electron microscopy images depict sarcomere structures of iPS-CMs in both control and stretch groups. **B** A comparison of sarcomere lengths between the control and stretch groups. The data are presented as means ± S.E.M., with n = 4 for the control group and n = 3 for the stretch group. Statistical analysis was conducted using an unpaired t-test. ****: P < 0.0001.

### 3.4. Evaluation of contractility of iPS-CMs

To compare contractile function of the iPS-CMs, we recorded video of the cultured cells under microscope. Spontaneous contraction of iPS-CMs was analyzed, and changes in contractility before and after stretching were visualized using vector field. An increase in contractility was observed in the stretch group (Figure 5A). Further, the change in the contractility before and after stretching was compared in both groups. Significant increase in contractility was observed in the stretch group compared to control group (−0.034 ± 0.061 and 0.411 ± 0.076, respectively) (Figure 5B). This indicates that the application of mechanical stretching during the induction of cardiomyocytes differentiation can improve the contractility of cardiac tissue.

**Figure 5.**
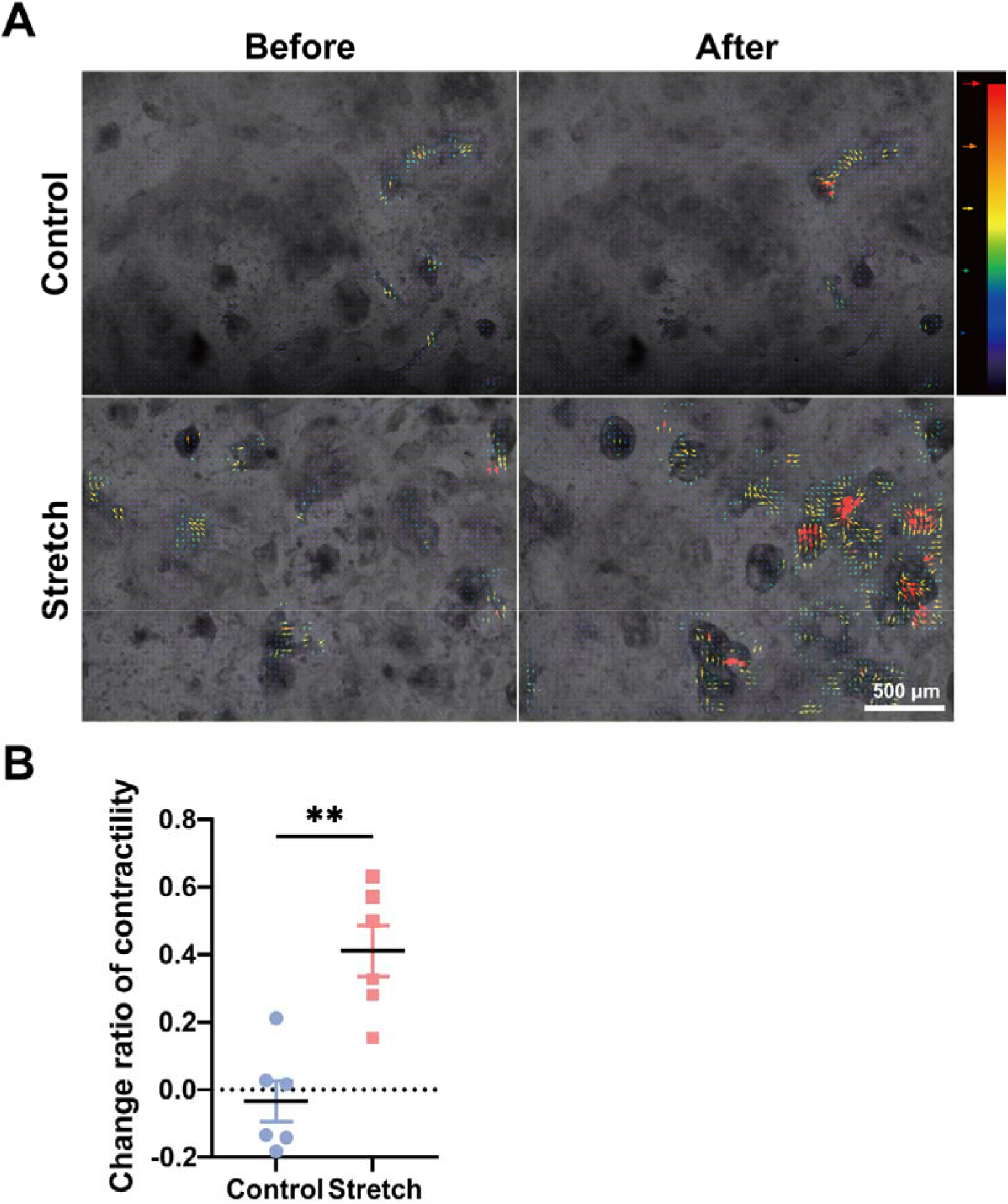
Contractility analysis of iPS-CMs. **A** Myocardial displacement vector field plot illustrating the direction and magnitude of displacement resulting from contraction, represented as vectors. **B** Comparison of the change ratio of contractility before and after stretching between the control and stretch groups. The data are presented as means ± S.E.M., with n = 6 for each condition. Statistical analysis was conducted using an unpaired t-test. **: P < 0.01.

## 4. Discussion

This experiment’s results show that the application of mechanical stretching stimulation in the process of inducing differentiation of iPS cells into myocardium can improve the maturity of iPS-CMs and increase the efficiency of differentiation.

The results of quantitative RT-PCR showed that compared with the control group, the myocardial marker cTnT and the mesodermal marker Nkx2.5 in the stretch group increased. Nkx2.5 can be found early in differentiation into cardiomyocytes, while cTnT is a marker in relatively mature myocardial tissue ^20^. The significant increase of cTnT and Nkx2.5 in the stretch group indicates that mechanical stretching stimulation promotes the differentiation of iPS cells into myocardial tissue. In addition, the decrease in pluripotent markers Oct4 and Sox2 is an indicator of the degree of pluripotent, indicating the differentiation efficiency of iPS cells. Compared with the control group, the expression of Oct4 and Sox2 tended to decrease in the stretch group, indicating that differentiation in the stretch group was enhanced.

Immunofluorescence staining of iPS-CMs results showed that cTnT-positive cells (red) and HCN4-positive cells (green) were observed in both groups, and they were high expressed in the stretch group compared with the control group. HCN4 channels are required for normal rhythm generation and conduction, playing a key role in cardiac pacing ^21^, and HCN4 overexpression shows enhanced I(F) current, spontaneous discharge, and pacing in iPS-CMs ^22^, which is consistent with our finding of increased cardiomyocytes contractility in the stretch group (Figure 5B). Experiments also confirmed the existence of sarcomere structure in the stretch group, these results indicating the maturation of myocardial tissue. Although HCN4 is closely associated with nodal cells, the transcript has also been detected in a high percentage of human primary atrial and ventricular cardiomyocytes, but at lower expression levels, such as in iPS-CMs ^23^. The results of previous publications evidenced that the pacemaker current I(F) present in primary ventricular cardiomyocytes ^24^. These data suggest that HCN4 is not specifically expressed in nodal cardiomyocytes. Indeed, using human embryonic stem cell-derived cardiomyocytes (hESC-CMs), Kim et al. showed that the presence of non-cardiomyocytes enhanced HCN4 ion channel development and electrophysiologic maturation. In early cardiomyocyte spheroid (CS) clusters, >99% of purified hESC-CMs stained positive for both cTnI (cardiac troponin I) and HCN4 staining ^25^. In Aoyama et al.’s experiments, the majority of iPS-CMs expressed the pacemaker marker HCN4 at early stages, however, expression declined in middle and late stages ^26^. Similarly, it was found that the majority of iPS-CMs identified as troponin T (TnT) positive cells were also positive for HCN4, but the pacemaker subtype had a higher expression than the contractile subtype at day 40, and with the downregulation of HCN4 expression in the later stages ^23^. As previously reported, this suggests that HCN4 is involved in the early maturation steps of cardiomyocytes ^27^. hESC-CMs have been reported to express HCN4 and have some pacing potency, but these potencies are lost with maturation ^28^. It is conceivable that HCN4 is retained only by the pacemaker cells when the cardiomyocytes tend to mature after appropriate conditions and long-term culture.

In order to observe the intracellular structure in more detail, a transmission electron microscope observation was performed, the arrangement of the sarcomere structure of the two groups was observed (Figure 4A). At the same time, in the stretch group, the greater arrangement of fibers in the myocardium was observed. It is known that the arrangement of fibers in healthy and mature cardiomyocytes is ordered, and studies have confirmed that changes in the arrangement of cardiomyocytes fibers affect the contractility of myocardial tissue. According to this report, cardiomyocytes with a normal arrangement will beat together, while cardiomyocytes with a disordered arrangement will lose the synchronized beating and various parts of the cell will beat in different directions ^29^. Therefore, the arrangement of fibers in cardiomyocytes is very important for producing myocardial tissue that exhibits highly synchronized beating.

When comparing the length of the sarcomere between the two groups, the control group was 1.232 ± 0.024 μm, while the stretch group showed a larger value of 1.537 ± 0.020 μm. The sarcomere length of mature myocardium is 1.6-2.2 μm ^30^, while the sarcomere length in the stretch group is closer to this. Therefore, the sarcomere structure can become more mature through mechanical stretching stimulation.

In the experiment, a video of spontaneous contraction was analyzed and recorded, and changes in contraction before and after stretching were visualized and quantified. As a result, an increase in contraction was observed in the stretch group. The experiment also compared the change rate of the average maximum contractility before and after stretching between the two groups. It was observed that the myocardial tissue in the control group increased contractility in some cases. On the other hand, all the samples showed no decrease of contractility in the stretch group, and an increase in the stability of contractility was observed. This indicates that the application of mechanical stretching stimulation during the induction of myocardial tissue differentiation can improve the contractility of myocardial tissue.

According to Matsuda et al., when iPS-CMs differentiation is induced by co-culture of HGF and iPS cells, maturation of iPS-CMs continues from the 15th day to the 30th day of culture ^10^. Therefore, this study decided to apply mechanical stretching stimulation from the 15th day of the culture. There are many papers on the stretching rate of stretching stimulation ^11,31^. According to Lux et al.’s research, under the stimulation of 2%, 5%, 10% and 20% stretching rate, when a mechanical stretching stimulation was applied with a 10% and 20% stretching rate to cardiomyocytes, the contractile function of cardiac structures was significantly reduced. And the application of 2% stretching did not induce any change in contractility compared to the control group ^32^. Also, 20% stretching rate induced cellular damage, leading to inflammation and apoptosis of cells ^33^. Therefore, although most mechanical stretching selected 1Hz to simulate human heart rate ^34,35,36,37^, in this experiment, under the conditions of co-culture with HGF, a mechanical stretching stimulation of 5% stretching rate and 0.5Hz frequency were applied during the differentiation of iPS cells into myocardial tissue, which is closer to in situ conditions of cardiomyocytes than normal culture dish conditions did not and induces cellular stress. In Lundy et al.’s experiment, Long-term culture of PSC-CMs increases cell size and promotes CMs maturation ^38^, however, it has been reported that with the progression of maturation, iPS-CMs become less responsive to mechanical stimulation ^39^. Considering the time and cost, more research is needed to explore the balance between them. Studies have shown that 48 hours of mechanical stretching stimulation is sufficient to induce cardiomyocyte maturation ^40,32^, in view of the 5% stretching rate and 0.5 Hz frequency is lower than the actual cardiac longitudinal strain (20.3 ± 5.6%) and heart rate (>1.0 Hz) ^41^, 72 hours of mechanical stretching stimulation was applied. Our research revealed that mature myocardial tissue can be obtained from iPS-HGF co-cultured CMs by mechanical stretching stimulation. It contributes to research in cardiac regenerative medicine and can provide a potential resource for application of iPS-CMs based on cardiac tissue repair.

## 5. Conflict of Interest

The authors declare that the research was conducted in the absence of any commercial or financial relationships that could be construed as a potential conflict of interest.

## 6. Author Contributions

YM, KT, HK and KN contributed to the conception and design of the study; MW and HI performed the experiments; MW, HI, CW, YIL, YUL wrote sections of the manuscript. All authors contributed to manuscript revision, and all authors read and approved the submitted version.

## 7. Acknowledgment

We express our gratitude to the Central Research Laboratory at Okayama University Medical School for their valuable assistance in preparation and observation of electron microscopy samples. This research received support from the Japan Society for the Promotion of Science (JSPS) through Grant-in-Aid for Scientific Research (B) (No. 20H04518) and Grant-in-Aid for Scientific Research (A) (No. 21H04960).

## Notes

### Competing Interest Statement

The authors have declared no competing interest.

